# Advanced PCL-Chitosan Nanofibrous Wound Care Material for Enhanced Wound Healing

**DOI:** 10.1101/2023.04.02.534760

**Authors:** Velikkakath G R, Jijo J Wilson, L V Thomas

## Abstract

Wet electrospinning has many advantages over conventional electrospinning processes. However, the technique is not commonly used due to the limited availability of suitable materials and accessibility of the technique. In this study different ratios of PCL-chitosan blend was prepared and fabricated into a highly porous scaffold system by wet electrospinning. The keratinocytes and fibroblast cells were seeded on the top and bottom side of the scaffold respectively to develop a bilayer skin construct. We conducted tests to examine the biological properties of the material including their cell viability, in vitro wound healing efficiency, and gene expression of collagen-I, collagen-III, cytokeratin-14, and cytokeratin-5. The findings indicate that PCL-chitosan can serve as a highly effective wound dressing material, with remarkable wound healing properties.

## Introduction

Skin is the largest organ in the human body and plays a crucial role in protecting the body from external harmful factors. Severe burn injuries, chronic wounds, and skin diseases can cause irreversible damage to the skin’s structure and function, leading to impaired healing and significant morbidity. The current clinical approaches for treating skin defects include autografts, allografts, and synthetic skin substitutes (1). However, these methods have several limitations, such as donor site morbidity, limited availability, and poor functionality (2).

Tissue engineering offers a promising alternative for repairing damaged skin by creating a functional skin substitute that mimics the natural skin structure and function. Electrospinning is a widely used technique in tissue engineering, producing micro-to nanofibers that mimic the natural extracellular matrix (ECM) and support cell adhesion, proliferation, and differentiation (3). Electrospun scaffolds can be tailored to match the desired mechanical, biochemical, and physical properties of the native skin, making them ideal for skin tissue engineering (4). However, due to the high fiber density and hydrophobicity, the electrospun scaffolds have limited cell penetration. Hence there is only a limited three-dimensional cell distribution happening in the electrospun scaffolds (5). Wet electrospinning is a modified version of electrospinning that involves the use of a liquid bath to collect the fibers. Since the fibers are collected in a liquid bath, the formed scaffold will have larger pores and less fiber density (6). This can be exploited for the development of a skin substitute or wound dressing for the treatment of dermal injuries.

The skin has a bilayer structure consisting of the epidermis and dermis. The epidermis consists of keratinocytes and the dermal layer carries the fibroblast cells. This bilayer structure can be easily recapitulated by wet electrospinning to make a skin substitute. Previous studies have shown scaffolds seeded with dermal cells, typically fibroblasts and keratinocytes, which are responsible for the production of collagen and other ECM components, favours the formation of the epidermis and dermis layer (7). Electrospun tissue engineered skin substitutes have been shown to closely mimic the structure and properties of natural skin, providing a favourable environment for cell growth and tissue regeneration (8).

One of the key advantages of electrospun tissue-engineered skin substitutes is their ability to be customized to meet specific patient needs. The scaffold can be engineered to mimic the thickness and architecture of the skin and can be designed to incorporate growth factors or other bioactive molecules to enhance tissue regeneration (9). The technology allows for the creation of nanofibrous scaffolds that closely mimic the structure and properties of natural skin, providing a favourable environment for cell growth and tissue regeneration. With further optimization and clinical testing, electrospun tissue engineered skin substitutes could become a viable option for the treatment of skin injuries and disorders (10). Electrospun tissue engineered skin substitutes have shown promising results in preclinical studies. The scaffolds have been shown to support cell migration, proliferation, and differentiation, and to promote the formation of a functional epidermis and dermis. Moreover, the electrospun skin substitutes have been shown to integrate with the host tissue and promote tissue regeneration (11).

Several electrospun tissue engineered skin substitutes are currently in various stages of development, with some already undergoing clinical trials. Some of these skin substitutes have shown positive results, with improved wound healing and tissue regeneration observed in patients with skin injuries and disorders. Among them, chitosan is widely used for the fabrication of wound dressing or as a skin substitute (12). However, due to the cationic nature and wide molecular weight distribution the material shows limited electrospinnability (13). There are very few studies that have shown the electrospinning of chitosan (14). For the ease of electrospinnability and to improve its mechanical strength chitosan can be blended with other polymers like polycaprolactone (PCL). PCL is known to be a bioinert polymer with attractive mechanical properties.

Chitosan is a natural biopolymer derived from chitin, which is found in the shells of crustaceans such as crabs, shrimps, and lobsters. Chitosan has been extensively investigated as a material for tissue engineering applications, including the development of skin substitutes. It has several properties that make it an attractive material for tissue engineering. It is biocompatible, biodegradable, non-toxic, and has antimicrobial properties. Moreover, it has been shown to support cell adhesion, proliferation, and differentiation, making it a suitable material for the fabrication of tissue engineered skin substitutes. In this study, polycaprolactone and chitosan were blended in different ratios and wet electrospun into a highly porous fibrous scaffold for skin tissue engineering.

## Materials and methods

### Wet electrospinning

The wet electrospinning was done using a customized electrospinning collector unit with an ethanol bath and a vertically placed nozzle. The PCL-chitosan blend of three different ratios was prepared using HFIP and acetic acid solvent. Ten percent PCL dissolved in glacial acetic acid and two percent chitosan dissolved in HFIP was blended at PCL: chitosan ratios of 3:7, 5:5, and 7:3. The electrospinning was performed at 13 kV voltage and a flow rate of 1 ml/ min. The nanofibers were collected in the ethanol bath placed at a distance of 7 cm from the nozzle.

### FTIR

The FTIR spectra of the samples were recorded using an FTIR spectrometer (BRUKER) in the ATR mode. The electrospun samples of the PCL-chitosan blend were used, and the spectrum was recorded in the 3500 - 500 cm^-1^ range using 25 scans, at a resolution of 4 cm^-1^.

### Mechanical test

The mechanical testing was done using solvent casted rectangular strips of 60 mm length, 12 mm width, and 0.02 mm thickness. The experiment was conducted using a load cell of 10 N and a crosshead speed of 5 mm/min in an instron UTM instrument (Universal Testing Machine).

### Contact angle measurement

The water contact angle was measured using a goniometer (Dataphysics) by the sessile drop method. On the surface of the sample, a drop (3 µl) of distilled water was placed using a glass syringe, and the contact angle was recorded within 3 seconds.

### Cell culture

NIH3T3 fibroblast cell lines and HaCaT keratinocyte cell lines were cultured using DMEM high glucose media (Gibco) containing 10% FBS (Gibco, South American grade) and 1x ABAM (Gibco). The cells were cultured at 37 °C with 5% CO_2_. The media was changed every two days.

### MTT assay

The MTT assay was performed as per ISO 10993-5. The HaCaT cells were seeded in 96 well plate at a density of 10000 cells/well and incubated overnight. The PCL-chitosan films were placed over the attached cells and cultured for 24 hrs in complete media. After culturing for 24 hrs, the media was removed and 50 µl of serum free media and 50 µl of MTT solution were added to the wells, the samples were then incubated for 3 hrs. After incubation 150 µl of DMSO was added. The absorbance was measured using a spectrophotometer at 590 nm.

### Scratch assay

For wound healing cell migration assays, HaCaT cells were seeded in 24 well plate at a seeding density of 0.5×10^6^ cells/ml. After reaching confluency, the cells were uniformly scraped vertically through the well using a sterile 200 µl pipette tip. After the wounding, the detached cells were washed off using PBS and media containing 10% FBS was added. The cells were continuously exposed to test samples (7 mm diameter and 1 mm thickness) for the indicated time periods. The scratched area was imaged every 12 hr using a phase contrast microscope (Olympus) at 10X magnification. The scratch area was analysed and quantified using imageJ.

### Immunostaining

The top side of the scaffold was seeded with HaCaT cells at a density of 5×10^5^ cells/ml. 3T3 fibroblast cells were seeded on the other side of the scaffold at a density of 5×10^5^ cells/ml. The cell-seeded scaffold was incubated for 90 min for the cells to attach. The construct was cultured for 2, 7, and 14 days in complete media containing 5ng/ml TGF-β (Thermofisher). The samples were retrieved after the respective culture period and fixed in 4% paraformaldehyde. After PBS wash, the samples were blocked with 3% BSA and incubated in the primary antibody (Vimentin - ab20346, Cytokeratin-14 - EPR17350) at 1:100 dilution at 4 °C overnight. The samples were washed thrice in PBS and incubated in 2° antibody at 1:2000 dilution (ab6800, ab150113) at room temperature for 1 hr. The nucleus was stained with DAPI at 1 nM concentration, for 15 mins. The images were captured using fluorescent microscope (Olympus).

### RT-PCR

The NIH3T3 and HaCat cells were cultured separately and the samples were harvested on 2, 7 and 14 days of culture. The RNA was isolated using TRIzol reagent (Thermofisher, USA). The total RNA concentration and ratio were measured using a NanoDrop (Thermoscientific, USA). Complementary DNAs were generated utilizing the PrimeScript RT reagent kit (Takara) and then amplified using RT-PCR with specific primers. PCR was conducted using a Real-Time PCR System (Analytik Jena, Germany), with each 20 μl of PCR mixture containing 10 μl of TB Green Premix Ex Taq II (Tli RNase H Plus, Takara), 0.4 μl of each PCR forward and reverse primers (10 μM), 0.4 μl of ROX reference dye (50×), 2 μl of template, and 6.8 μl of sterile purified water. Samples were subjected to cycles of incubation at 95 °C for 30 s, 40 cycles at 95 °C for 5 s and 60 °C for 34 s, and a final cycle at 95 °C for 15 s, 60 °C for 1 min, and 95 °C for 15 s. The outcomes were assessed using the comparative cycle threshold (ΔΔCT) method to determine gene expression fold changes that were standardized to the levels of GAPDH gene transcripts. The RT-PCR was performed using the primers sequences given in Table-1.

**Table-1.**
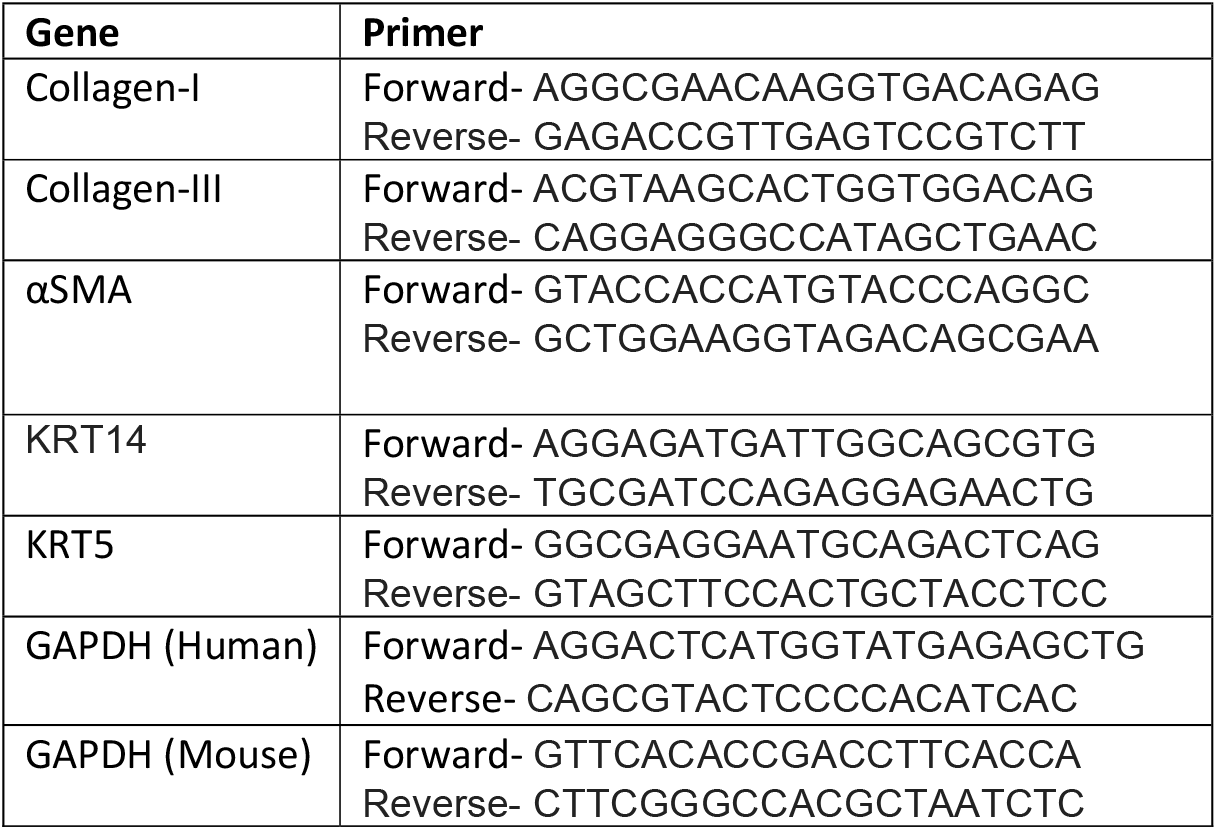

**Table-2.**

| Sample | Contact angle (°) |
| --- | --- |
| PCH37 | 90±4.6 |
| PCH55 | 112±7.3 |
| PCH73 | 118±6.7 |

## Results

### SEM analysis

A major limitation of electrospun scaffolds is their high fiber density and lack of cell penetration. In this study, the PCL-chitosan material is electrospun into an ethanol bath to fabricate scaffolds having loosely packed fibers with larger pore sizes than the conventional electrospun scaffolds. The morphology of the electrospun samples was studied by SEM microscopy. Figure 1 shows the SEM images of the conventionally electrospun and wet electrospun PCL-chitosan scaffolds. (Figure 1) Shows, in the conventionally electrospun scaffolds, the fiber diameter increased with the increase in the chitosan content, and the fibers were tightly packed having pore sizes of 0.242 µm±0.84 µm, 1.4±1.1 µm, 2.4 µm±0.432 µm for PCH37 (Figure 1 A), PCH55 (Figure 1 B), PCH73 (Figure 1 C) respectively. Whereas the wet electrospun system showed larger pore sizes of 84 µm±26 µm, 72 µm±18 µm, and 93 µm±24 µm for PCH37, PCH55, and PCH73 respectively. Unlike the conventionally electrospun samples, the wet electrospun samples did not show a change in pore size with the change in the chitosan ratio.

**Figure 1:**
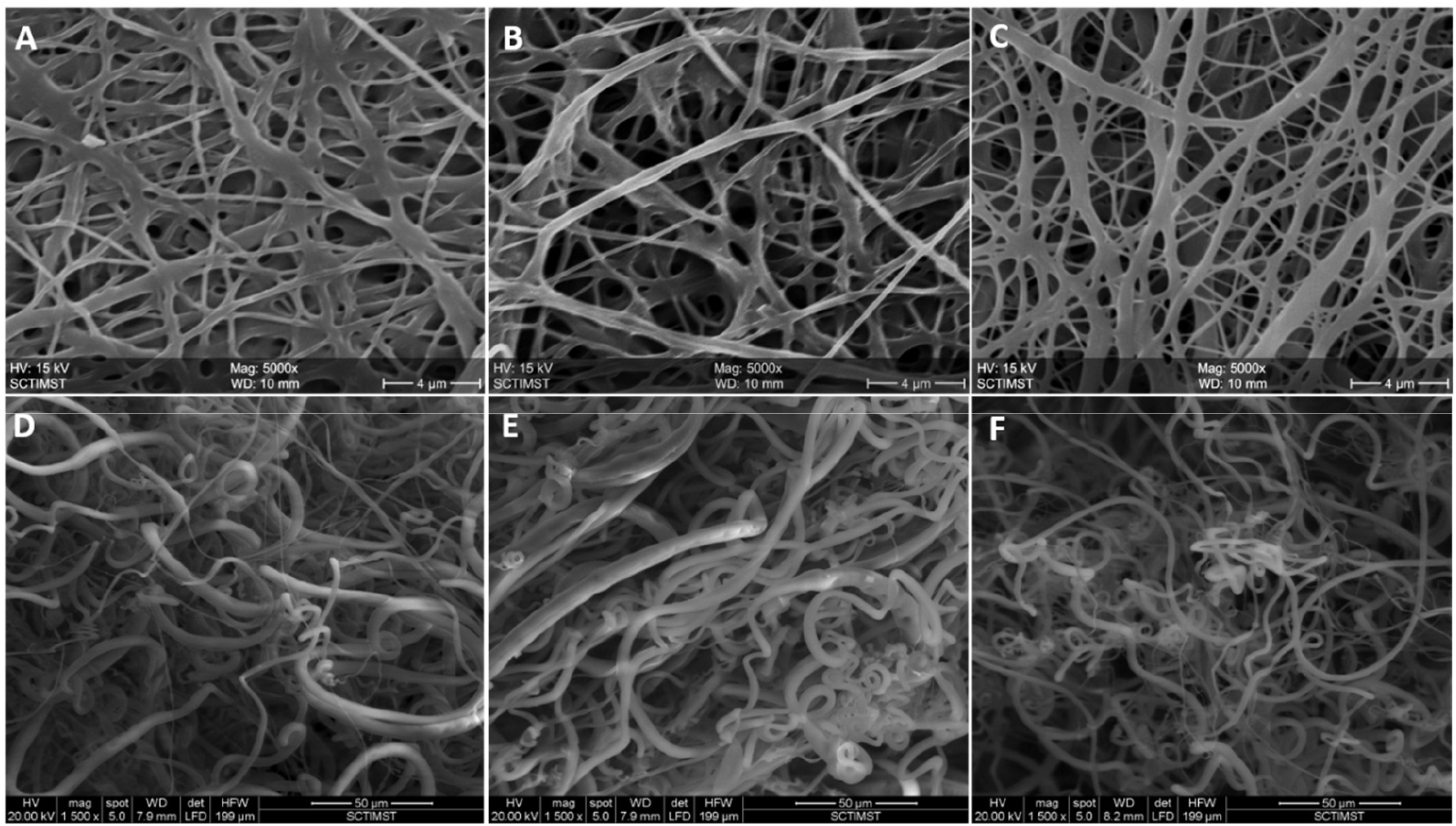
SEM images showing PCH scaffold prepared by conventional electrospinning (A, B, C) and wet electrospinning (D, E, F).

### FTIR analysis

The chemical structure of the PCL-chitosan blend was confirmed by FTIR spectroscopy. Figure 2 shows the FTIR spectrum of the three different ratios of PCL-CH blends. Peaks at 1726 cm^-1^, 1245 cm^-1^, 890 cm^-1,^ and 1117 cm^-1^ are characteristic PCL peaks that are related to stretching of the C-O and C-O-C bridges, CH_2_ deformation, and carbonyl ester bonds respectively. The peaks at 897 cm^-1^, 1080 cm^-1^, 1189 cm^-1^, 1485 cm^-1,^ and 2930 cm^-1^ are attributed to the CH_3_OH, C-O stretching, asymmetric stretching of the C-O-C bridge, N-H bending, and C-H stretching respectively of chitosan. The FTIR of PCH electrospun nanofiber contains the typical PCL and chitosan (CH) peaks.

**Figure 2:**
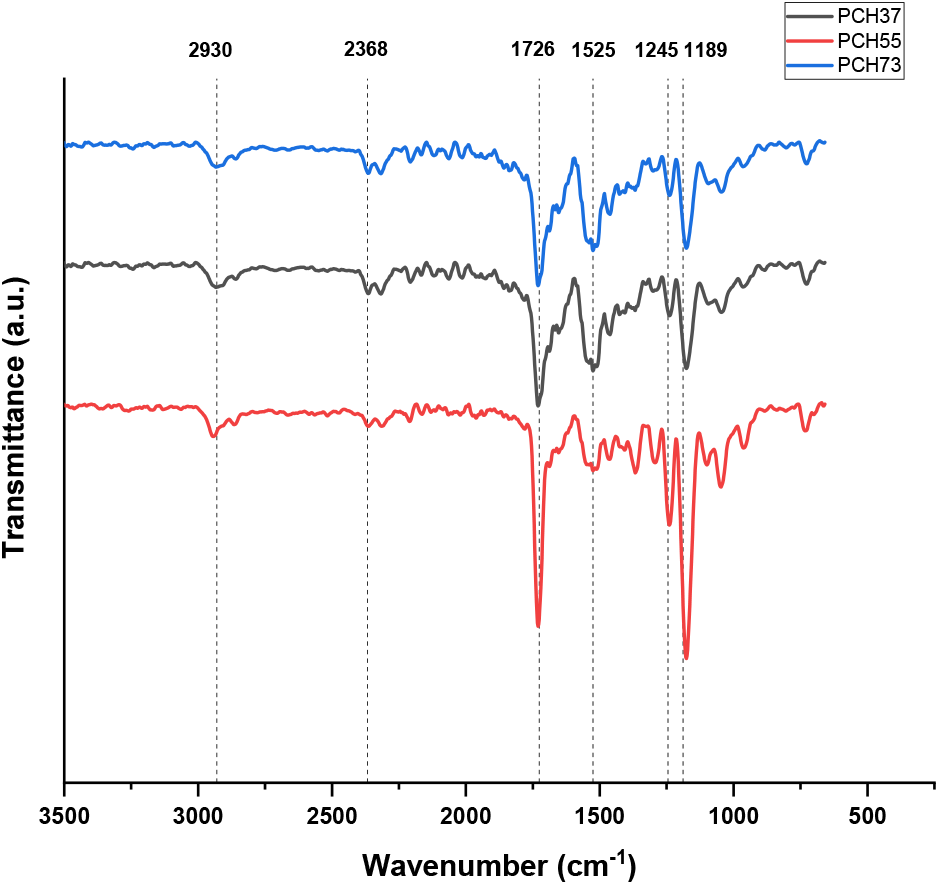
FTIR spectrum of PCH37, PCH55, and PCH73 showing the characteristic peaks of chitosan and PCL.

### Mechanical testing of PCL-chitosan blend

For use in wound dressing applications, the material needs to have the right mechanical characteristics, particularly elongation at break. It means that a wound dressing must fulfill specific requirements in order to properly cover a wound and prevent rupture as the wound heals. Tensile stress-strain curves for the PCL-chitosan blend films of different ratios are shown in Figure 3. The material’s tensile strength ranged from 5.6±1.4 MPa to 8.7±3.1 MPa, which is suitable for use as a wound dressing material. The lowest tensile strength was exhibited by the PCH73 sample and the highest tensile strength by the PCH37 sample, which shows the tensile strength increases with the addition of chitosan. However, the elongation at break showed an opposite trend, the lowest elongation at break of 10% was shown by PCH37 and PCH73 showed the highest value of 140%. Hence, the elastic property of the material increases with an increase in the PCL content.

**Figure 3:**
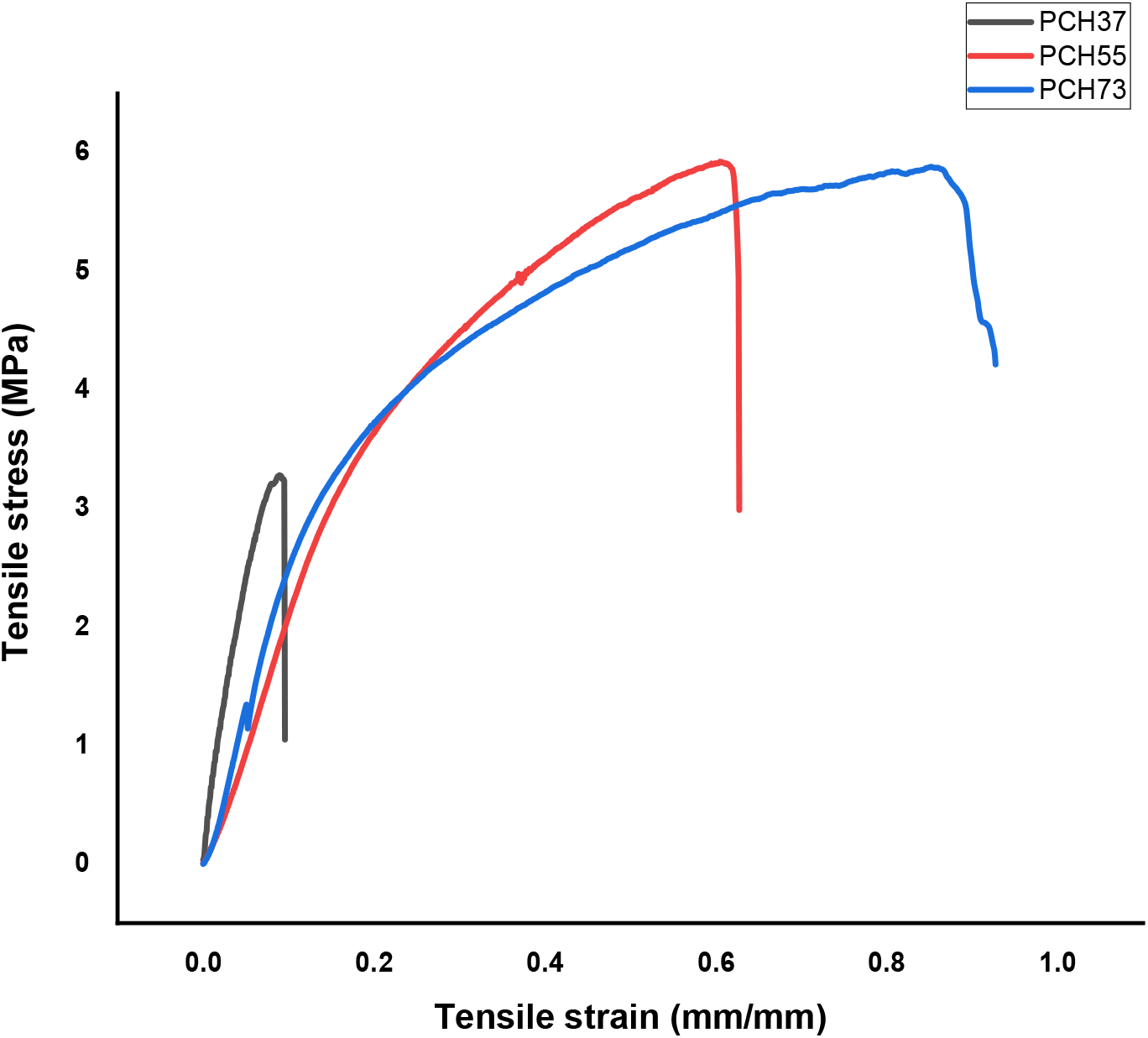
Stress-strain curve of PCH showing an increase in the elasticity with an increase in the PCL content.

### Water Contact angle measurement of PCL-chitosan blend

The in vivo interplay between the surface of the damaged tissue and the wound dressing is revealed by the contact angle between the water and the wound dressing. Increased cell adhesion and proliferation are generally facilitated and enhanced by increasing the hydrophilicity of the wound dressing surface. Table 1 lists the electrospun nanofibers’ water contact angle (WCA) values. Due to PCL’s hydrophobicity, the contact angle of the material ranged between 90±2.6° - 118±6.7°. Due to the amino ester and hydroxyl groups in the CH chains, the addition of CH increased the surface hydrophilicity of PCL (Figure4).

**Figure 4:**
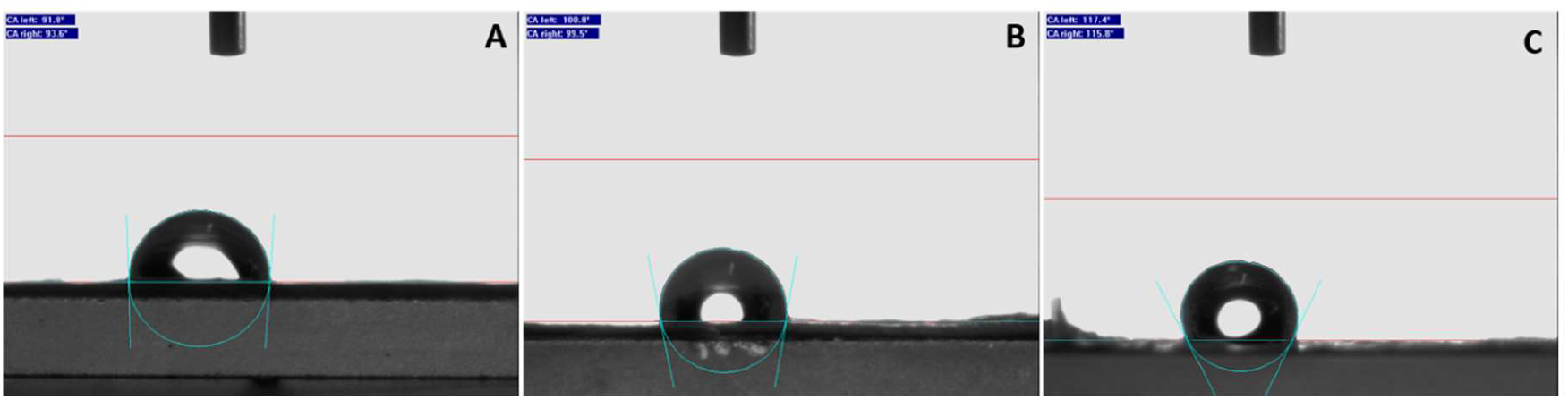
Contact angle measurement of (A) PCH37, (B) PCH55, PCH73 ratios, showing an increase in hydrophobicity with an increase in PCL content.

### MTT Assay showed the prepared material is non-cytotoxic

Assessing cell viability is crucial for determining the potential cytotoxicity of the material as a skin substitute or wound dressings. To evaluate this, the viability of HaCaT cells was measured over a period of 24 hrs in the presence of different ratios of prepared PCH. The viability of the cells was determined by measuring cell proliferation by MTT assay. The viability of cells cultured in presence of PCH37, PCH55, PCH73, and PCL was similar to the tissue culture control and showed above 90% cell viability in all the ratios (Figure 5). This indicates that the material is non-cytotoxic and does affect the normal growth and proliferation of the cells.

**Figure 5:**
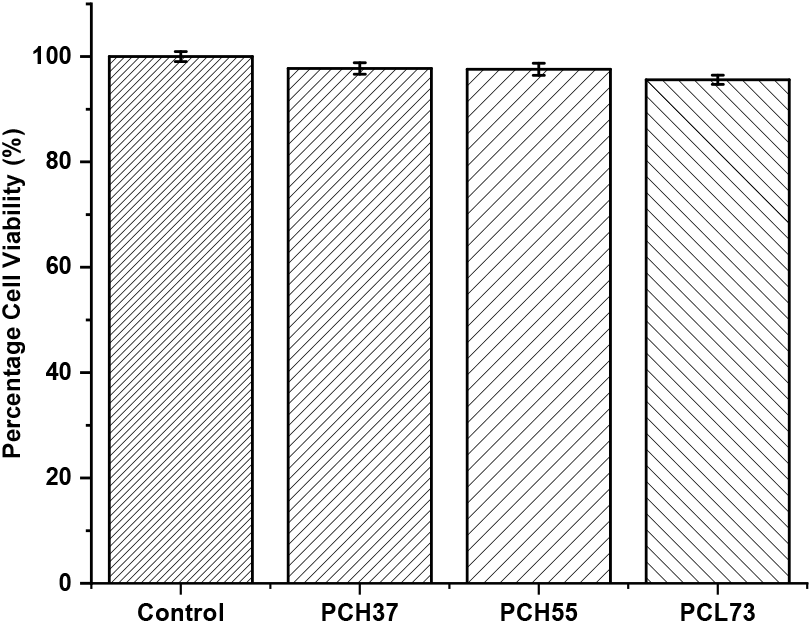
MTT assay showing the 24 hrs cell viability of PCH37, PCH55, and PCH73 samples.

### Scratch assay showed the addition of chitosan improved wound closure

Keratinocytes are a crucial component of the epidermis that play a vital role in maintaining the skin barrier and repairing skin injuries through epithelialization. Re-epithelialization is a crucial stage in the wound healing process, which involves increased migration and proliferation of keratinocytes over the wound area. The results of the scratch assay are presented in Figure 5, 6. The samples treated with PCH37, PCH55, and PCH73 showed an increase in wound healing activity. However, in case of PCH37 ratio, the wound healing was faster compared to the other groups. The cells started migrating after 12 hours of culture and 80% of the wound was covered in 48 hrs of culture in the PCH37 sample. In all the time points PCH37 showed the highest wound closure compared to the other ratios.

**Figure 6:**
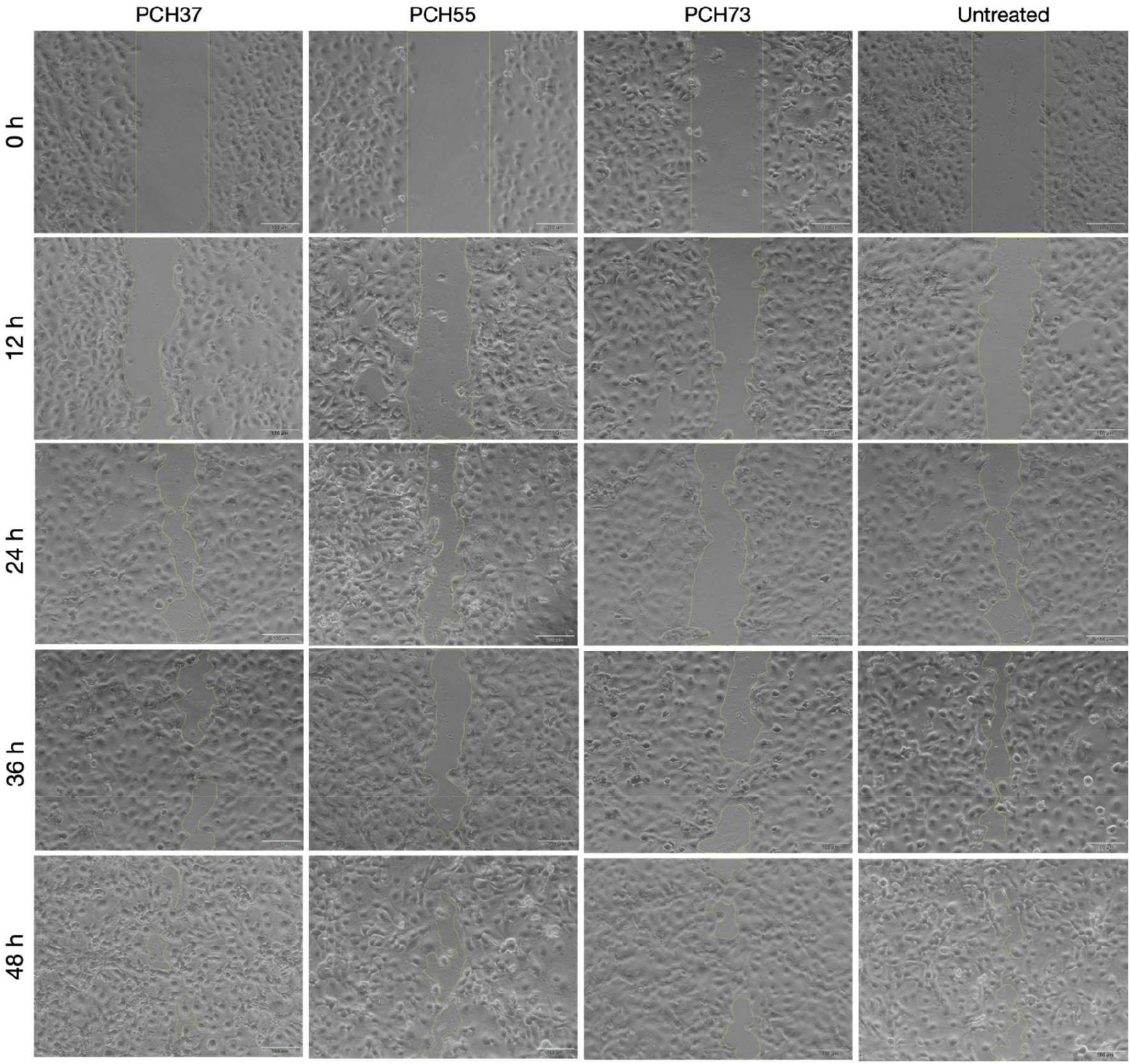
The scratch test of (A) PCH37, (B) PCH55, (C) PCH 73 showing wound closure.

**Figure 7:**
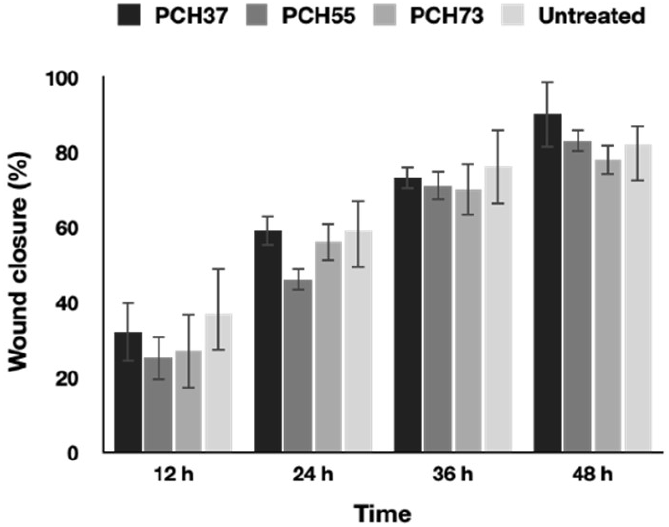
Area measurement showing PCH37 with maximum wound closure in 48 hours.

### Immunostaining

Since the PCH37 showed superior wound closure and cell viability compared to the other ratios, this ratio was selected for further studies. The bilayer construct generated by seeding keratinocytes and fibroblasts was characterized by evaluating the expression of their respective markers. The keratinocytes were stained positive for cytokeratin-14 and negative for vimentin. Whereas, the fibroblast cells expressed vimentin but not cytokeratin-14. Due to the highly porous structure of the PCH37 scaffold, the cells condensed and formed a cluster (Figure 8). Thus, the 3D fibrous scaffold system was favouring cell migration and formation of a 3D tissue without affecting the functionality of the cells.

**Figure 8:**
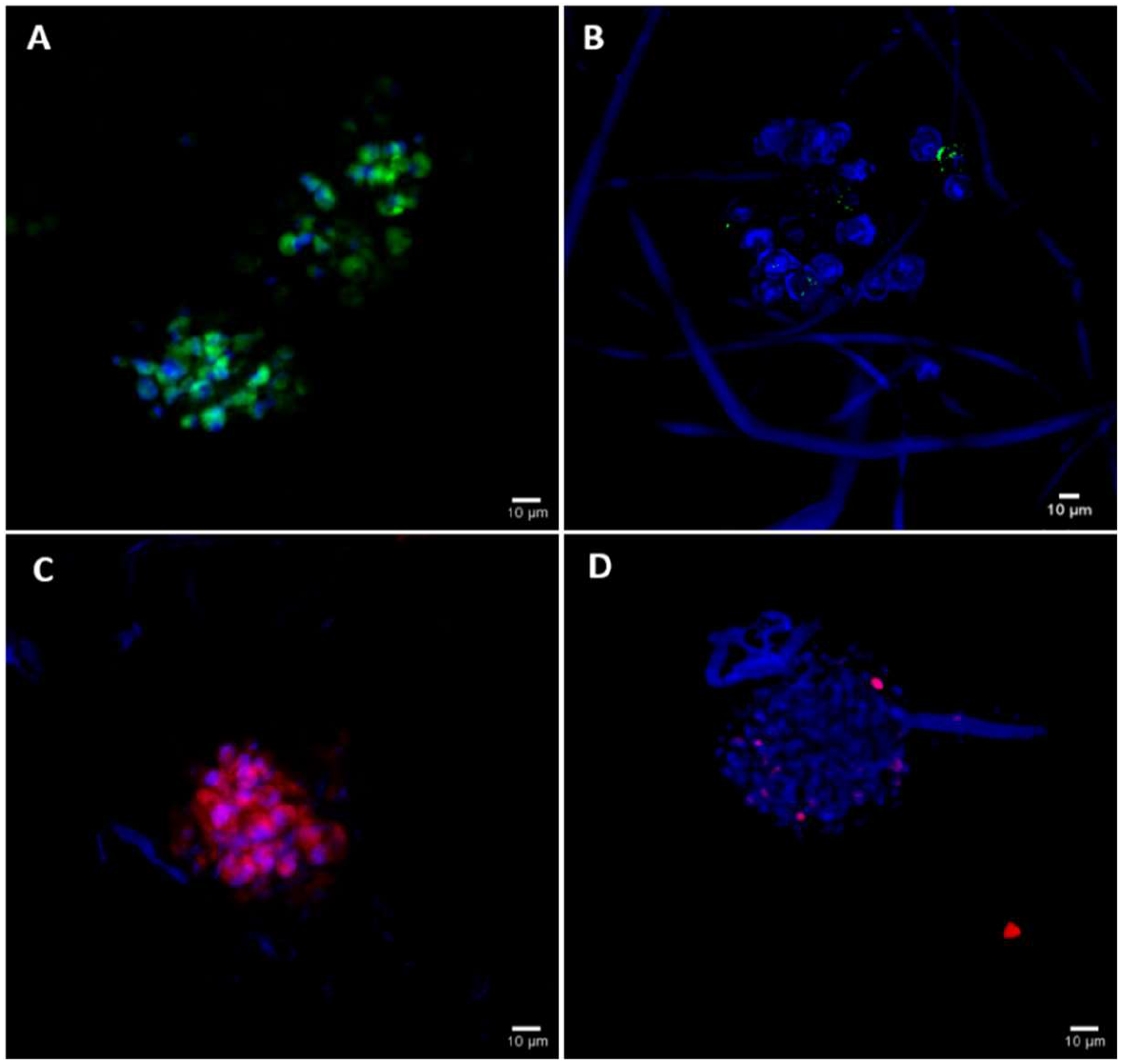
Expression of marker proteins of fibroblast and keratinocyte cells after 7 days of culture in PCH37 scaffold. (A) Vimentin expression in fibroblast cells, (B) Cytokeratin-14 expression in fibroblast cells, (C) Cytokeratin-14 expression in keratinocytes, (D) Vimentin expression in keratinocyte cells.

### Gene expression

The study used RT-PCR analysis to examine the expression of collagen type I, III, αSMA, KRT5, and 14 genes (Figure 9). The ECM proteins collagen I and III play a major role in the remodelling of the dermis. The fibroblast cells seeded on the scaffold showed expression of collagen-I only in the later time points. Whereas the collagen-III was found to be expressed from 2 days of the culture. The col-I showed expression from the 7^th^ day of the culture and showed an 8.8-fold increase by the 14^th^ day of the culture. Unlike col-I, col-III showed maximum expression on the 7^th^ day of culture and its expression was found to be decreasing by the 14^th^ day. The cytoskeletal protein αSMA was found to be downregulated in the 7 days and 14 days of culture, probably due to the formation of clusters in the scaffold. The keratinocyte cytoskeletal filament KRT5 and 14 showed similar expression in all the time points.

**Figure 9:**
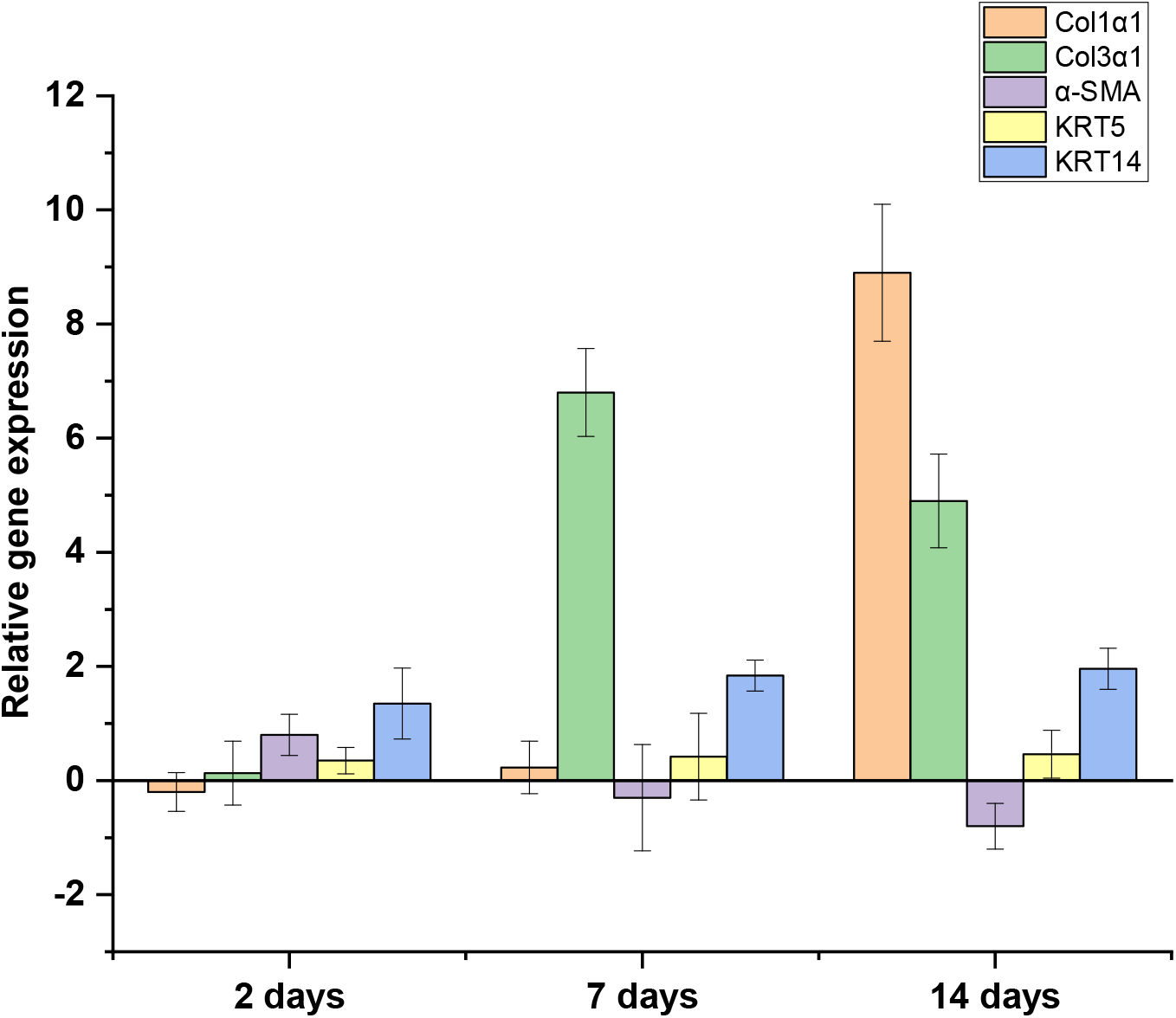
RT-PCR analysis of PCH37 scaffold seeded with keratinocyte and fibroblast cells, showing expression of ECM and cytoskeletal proteins.

## Discussion

In this study, we were able to create a wound dressing made using a combination of PCL and chitosan by wet electrospinning. This dressing was designed to provide better cell infiltration. Moreover. The presence of chitosan is expected to confer anti-bacterial and hemostatic properties to the system. The dressing’s highly porous structure encourages free movement and growth of keratinocytes and fibroblasts.

In recent years, there has been a growing interest in using electrospun PCL meshes for skin regeneration and wound dressing. Various types of functionalized PCL have been utilized to create electrospun fiber meshes with different structures for wound healing applications (15). Thus, recent studies have focused on using PCL-based blends and composites, or modifying electrospun fiber meshes to improve their physical, chemical, and bioactive properties. For instance, researchers have found that incorporating fibroblasts into electrospun fibers made from PCL-collagen blend with chitosan nanoparticles, can significantly enhance wound healing compared to using cell-free membranes or control treatments. The wounds treated with cell-loaded PCL-collagen fibers with chitosan nanoparticles showed faster healing than those treated with pure PCL meshes or control, with complete healing observed after 14 days (16). These findings, corroborates our in vitro wound healing results, the early wound closure observed in the PCH37 ratio could be because of the increased ratio of chitosan. Moreover, the immunostaining results showed, the mesenchymal origin marker vimentin is not expressed in keratinocytes and was found to be expressing in fibroblast cells. And cytokeratin-14, a marker of basal keratinocytes was expressed only in the keratinocytes. This confirms that the cells seeded on the scaffold are not dedifferentiating and are maintaining its functionality.

When it comes to wound dressing materials, it is important that they are easy to handle and apply. Mechanical testing can be used to assess this ability. The tensile strength of PCL-CH ranges from 5.6±1.4 MPa to 8.7±3.1 MPa, indicating good mechanical strength for wound dressing applications. The porosity of the dressing is also important, as it affects both the mechanical properties and water retention capacity. Porous structures can help with cell attachment, migration, and expansion by easily absorbing culture medium. The chitosan has improved the hydrophilicity of the PCL, which is important for cell adhesion and growth. However, the dressing shouldn’t be too wet or too dry, as this can negatively impact wound healing. Biodegradable synthetic polymers tend to be hydrophobic, but adding chitosan can improve wettability. The larger pore size can improve the free volume available for water molecule transfer. This will prevent the material from drying and improve the cell infiltration.

The process of wound healing is typically comprised of four stages, which are referred to as hemostasis, inflammation, proliferation, and skin remodelling (17). Chitosan and its derivatives have a significant impact on the first three stages of wound healing. In the hemostasis stage, they aid in stopping bleeding by promoting platelet and erythrocyte aggregation and preventing the breakdown of fibrin (18). During the inflammation stage, they assist in removing bacteria from the wound (19). Finally, in the proliferation stage, they accelerate skin growth by promoting the formation of granulation tissue (20).

After the inflammatory stage the granulation tissue formed has the ability to replenish the damaged region and facilitate the regeneration of the epidermis. Studies have revealed that chitosan can expedite the healing process of skin wounds by encouraging the growth of inflammatory cells like macrophages, fibroblasts, and blood vessels. Chitosan can stimulate macrophages to release cytokines, including transforming growth factor-β (TGF-β), PDGF, and IL-1. TGF-β stimulates the migration of macrophages to injured regions, encourages the proliferation of fibroblasts, and increases the secretion of collagen (21). This was evident in the gene expression data as well. Even though collagen-I was absent in the early time points, by 14^th^ day there was prominent expression of collagen-I. Whereas, collagen-III is expressed in the early stages of wound healing and gets replaced by the collagen-I, this could be the reason for the expression of collagen-III in the early time points and decreases by 14^th^ day. Thus our in vitro results show that, the fabricated wound dressing could be a potential candidate for the clinical studies to improve the wound healing.

## Conclusion

Electrospinning is a versatile and promising technique for producing tissue-engineered skin substitutes. Electrospun scaffolds can be tailored to match the mechanical, biochemical, and physical properties of the native skin and support the attachment, proliferation, and differentiation of skin cells. The electrospun tissue-engineered skin holds great potential for treating skin defects and improving the quality of the life of patients with severe skin injuries or diseases. However, further studies in the animals are required to validate the efficiency of the device in the in vivo system.

## Acknowledgement

The authors thank the Director, SCTIMST, and Head, BMT wing, SCTIMST, for providing the facilities and infrastructure to carry out this work.

## Notes

### Competing Interest Statement

The authors have declared no competing interest.

